# A2A Receptor contributes to tumor progression in P2X7 null mice

**DOI:** 10.1101/2022.02.28.482298

**Authors:** Elena De Marchi, Anna Pegoraro, Roberta Turiello, Francesco Di Virgilio, Silvana Morello, Elena Adinolfi

**Author notes:** **Correspondence:** Elena Adinolfi.

## Abstract

ATP and adenosine are key constituents of the tumor niche where they exert opposite and complementary roles. ATP promotes tumor growth but also immune eradicating responses mainly via the P2X7 receptor (P2X7R), while adenosine acts as a potent immune suppressor and facilitates neovascularization thanks to A2A receptor (A2AR) activity. However, studies exploring the interplay between P2X7R and A2AR in the tumor microenvironment are as yet missing. Here we investigated tumor growth in C57/bl6 P2X7 null mice inoculated with B16-F10 melanoma cells, showing that several pro-inflammatory cytokines (IL1-β, TNF-α, IL-6, IL-12, IL-17, IFN-γ) were significantly decreased while the immune suppressant TGF-β was almost three-fold increased. Interestingly, tumors growing in P2X7-null mice also upregulated tumor-associated and splenic A2AR, suggesting that immunosuppression associated to lack of the P2X7R might depend upon A2AR overexpression. Immunohistochemical analysis showed that tumor cells A2AR expression was increased, especially around necrotic areas, and that VEGF and the endothelial marker CD31 were also upregulated. The A2AR antagonist SCH-58261 reduced tumor growth similarly in the P2X7 WT, or null mice strain. However, SCH-58261only reduced VEGF in the P2X7-KO mice, thus supporting the hypothesis of an A2AR mediated increase in vascularisation in P2X7-null host. SCH-58261 administration also significantly reduced intratumor TGF-β, thus supporting a key immune suppressive role of A2AR in our model. This study shows a novel direct correlation between P2X7R and A2AR in oncogenesis and paves the way for new combined therapies promoting anti-cancer immune responses and reducing tumor vascularization.

## Introduction

The P2X7 receptor (P2X7R) is an ATP gated ion channel known to be central in inflammation for its ability to activate the NLRP3 inflammasome and trigger IL-1β release (Di Virgilio et al., 2017; Orioli et al., 2017; Adinolfi et al., 2018; Pelegrin, 2021). In the context of cancer, P2X7R has been assigned multiple, and often contrasting, roles as a driver of cancer cell growth (Di Virgilio et al., 2018; Scarpellino et al., 2019) and metastatic dissemination (Di Virgilio et al., 2016; Ulrich et al., 2018; Pegoraro et al., 2021), or as a promoter of immune mediated tumor eradication (Adinolfi et al., 2015; Adinolfi et al., 2019; De Marchi et al., 2019; Lara et al., 2020). Interestingly, approaches based on either P2X7R antagonism or agonism have proved effective in reducing tumor growth, thus leaving many open questions on the mechanisms underlying P2X7R activity in cancer (Adinolfi et al., 2012; Amoroso et al., 2015; Douguet et al., 2021; Drill et al., 2021). The abundance of P2X7R ligand extracellular ATP (eATP) is a well-recognized characteristic of the tumor microenvironment (TME) (Di Virgilio et al., 2018; De Marchi et al., 2020). The TME is also rich in the ATP hydrolysis derivative extracellular adenosine (eADO) (Boison and Yegutkin, 2019). These two molecules exert opposing actions on the immune system, as, while eATP (via P2X7R) is pro-inflammatory and promotes anti-tumor immune response (Kepp et al., 2021), adenosine acts as an immunosuppressant, thus facilitating tumor immune escape (Leone and Emens, 2018; Young et al., 2018; Allard et al., 2020; Antonioli et al., 2021). However, the effects of both eATP and eADO are not limited to activity on immune cells as often through the same receptors expressed by either tumor cells or surrounding stroma they also promote cancer growth, vascularization and metastasis (Di Virgilio et al., 2016; Borea et al., 2018; Scarpellino et al., 2019; Arnaud-Sampaio et al., 2020; Sitkovsky, 2020).

In the TME, eADO accumulation causes suppression of immune effector functions and promotes tumor progression (Allard et al., 2020). These effects are mediated via A2A and A2B receptors (A2AR, A2BR), which are expressed by immune stromal and tumor cells. These receptors are both coupled to Gs protein and differ for their ligand binding affinity as the A2AR has a high (Kd: 310 nM), while A2BR has a low affinity (Kd 15 μM) (Yegutkin, 2008). The A2AR has been widely studied, emerging as a crucial mediator of eADO effects in the TME (Ohta et al., 2006; Boison and Yegutkin, 2019). Its activation potently suppresses both B and T cells’ effector activity while promoting the accumulation of regulatory T cells (Tregs), which are known to be immunosuppressive (Sorrentino et al., 2013; Antonioli et al., 2021). Furthermore, stimulation of A2AR, on macrophages populating the tumor lesion, promotes the differentiation of M2-like macrophages, which are the major responsibles for the accumulation of vascular endothelial growth factor (VEGF) in the TME and the subsequential angiogenesis (Sitkovsky et al., 2014; Allard et al., 2020). Importantly, A2AR not only promotes the establishment of a pro-tumorigenic environment driving macrophage differentiation, but also mediates the direct effects of eADO on tumor cells, as shown by the finding that in animal models, tumor cell A2AR promotes invasiveness, motility and proliferation (Beavis et al., 2013; Koszalka et al., 2016; Allard et al., 2019; Kamai et al., 2021).

The eATP and eADO levels are strictly intertwined. In the extracellular milieu, eADO is generated starting from eATP via two ectoenzymes, CD39 and CD73, which are also upregulated in the tumor lesion, favouring cancer growth through eADO-mediated immune suppression (Allard et al., 2020; Battastini et al., 2021). We have recently demonstrated that, in implanted murine melanoma, P2X7R genetic knockdown or pharmacological antagonism modulates CD39 and CD73 levels on Tregs, CD4^+^ effector lymphocytes, macrophage and dendritic cells (De Marchi et al., 2019). Changes in ectonucleotidases expression correlate with alterations in eATP levels (De Marchi et al., 2019), strongly suggesting that P2X7R expression and function affect the entire eATP/eADO axis in the TME. Independent studies also indicated a key role of the P2X7R in anti-tumor, CD39-targeting therapies (Li et al., 2019; Yan et al., 2020). Despite this wealth of studies, to our knowledge, no studies demonstrating a correlation between P2X7R and A2AR activity in cancer have been carried out so far. Here we show that the A2AR is upregulated in melanoma bearing P2X7R-null mice and highlight a pivotal role for the A2AR in immune suppression and neovascularization promoted by the absence of the P2X7R. We thus demonstrate, for the first time, a P2X7R-A2AR centred crosstalk in tumorigenesis.

## Materials and Methods

### Cell cultures

Mouse B16-F10 (ATCC CRL-6475) melanoma cell lines were purchased from Sigma Aldrich and periodically tested with MycoAlertTM kit (Lonza, Switzerland). B16-10 were grown in RPMI-1640 medium (Sigma-Aldrich) plus non-essential amino acids (Sigma-Aldrich), FBS (10%), penicillin (100 U/ml), and streptomycin (100 mg/ml) all purchased from Euroclone, Milan, Italy.

### Experiments in murine models

In vivo experiments were performed with a *p2x7^-/-^* mice in the C57bl/6 strain and corresponding wild type controls (Adinolfi et al., 2015; De Marchi et al., 2019). Based on calculations performed with the G-power software (Faul et al., 2007) on previously published data (De Marchi et al., 2019), a minimal sample size of 6 animals per group was chosen to achieve a predicted power of 0,85 with an effect size of 1,78 using a two-tailed t-test. B16-F10 cells (2,5 × 10^5^) were subcutaneously injected into the right flank of each mouse. The animals were randomised, thanks to the free software available at random.org, and the operator was blinded to the group of allocation. Tumors were measured with a calliper, and their volume calculated according to the following equation: volume = π/6 [w1 × (w2)^2^], were w1 = major diameter and w2 = minor diameter. A2AR antagonist SCH-58261 (Tocris, bio-techne) or placebo (sterile PBS containing 0.005% DMSO) were intra-peritoneum injected (100 μl) every three days after first tumor mass detection (day five from inoculum). SCH-58261 dose (1 mg/kg) and administration schedule were similar to those previously described in literature (Young et al., 2016; Ma et al., 2017). Mice blood samples were collected from the submandibular vein and complemented with 1,5 mg/ml EDTA, immediately before sacrifice (post inoculum day 14). Tumors were post-mortem excised and processed for further analysis. Mice plasma was collected by centrifugation of blood (1000 x g, 10 minutes at 4°C). Halt Protease and Phosphatase Inhibitor Cocktail (Thermo Scientific) was added to the samples before storage at −80°C. All animal procedures were approved by the Organism for the animal wellbeing (OPBA) of the University of Ferrara and the Italian Ministry of Health in compliance with EU Directive 2010/63/EU and Italian D.Lgs 26/2014.

### Cytokines and growth factors evaluation

Cytokines levels were evaluated within diluted plasma (1:2) either with mouse interleukin (IL)-1β, tumor necrosis factor (TNF)-α, transforming growth factor (TGF)-β1 (Boster, distributed by Tema Ricerca, Bologna, Italy), interferon (IFN)-γ (R&D, bio-techne) ELISA kits or with Luminex mouse multi-cytokine assay kit (R&D, bio-techne) as per manufacturer’s instructions. Tumor mass were homogenised in lysis buffer (300 μM sucrose, 1 mM K2HPO_4_, 1 mM MgSO_4_, 5.5 mM glucose, 20 mM HEPES (pH 7.4), Halt™ Protease and Phosphatase inhibitor cocktail, EDTA-free 100× (Thermo Fisher), IGEPAL CA-630 0.5%) and loaded for VEGF and TGF-β1 analysis in a mass ratio of 1:5 with mouse ELISA kits (Boster, distributed by Tema Ricerca, Bologna, Italy).

### Immunoblotting

Tumor or organs were homogenised in lysis buffer (300 μM sucrose, 1 mM K_2_HPO_4_, 1 mM MgSO_4_, 5.5 mM glucose, 20 mM HEPES (pH 7.4), Halt™ Protease and Phosphatase inhibitor cocktail, EDTA-free 100× (Thermo Fisher), IGEPAL CA-630 0.5%). The lysates were loaded in 4–12% NuPAGE Bis-Tris precast gels (Thermo Scientific). Following electrophoretic separation we perfomed transfer blotting on nitrocellulose membranes (Amersham Protran, GE Healthcare, USA); which were subsequently incubated overnight at 4 °C with primary antibodies as follows: anti-P2X7 antibody (P8232, Sigma-Aldrich) was diluted 1:300, anti-A2AR antibody (sc-32261 Santa Cruz Biotechnology, Heidelberg, Germany) was diluted 1:2500, anti-myosin II antibody (M8064 Sigma) was diluted 1:1000, anti-GAPDH antibody (#6C5-sc32233, Santa Cruz Biotechnology) was diluted 1:1000 and anti-β actin antibody (#E-ab-20031, Santa Cruz Biotechnology) was diluted 1:2500. Goat anti-rabbit (170-6515, BIORAD) or goat anti-mouse (170-6516, BIORAD), HRP-conjugated antibodies were applied at a 1:3000 dilution as secondary antibodies. Protein bands were visualised by ECL HRP Chemiluminescent Substrate ETA C ULTRA 2.0 (Cyanagen Srl, Bologna, Italy) with a Licor C-Digit Model 3600. Densitometric analysis was carried out with ImageJ software, and data were normalised on the appropriate housekeeping protein (myosin II, GAPDH or β actin).

### Immunohistochemistry

Excised tumors were fixed in Bouin (Sigma Aldrich) for 7 hours at 4°C, dehydrated in cold-graded ethanol series, cleared in xylene, and embedded in paraffin. For immunohistochemistry of A2AR and CD31, serial 7-μm-thick sections were rehydrated, washed in TBS (150 mM NaCl, 50 mM Tris, pH 7.6), incubated for 1 hour in blocking solution (TBS 8% FBS, 5% BSA 0,2% TRITON X-100) and subsequently for 16 hours at 4°C in blocking solution containing primary antibodies (anti-A2AR mouse antibody, sc-32261 Santa Cruz Biotechnology, at 1:100 and anti-CD31 rat monoclonal antibody, ER-MP12 Thermo Scientific, 1:20). For detection of A2AR and CD31, heat-induced epitope retrieval was performed using Rodent Decloaker (RD913M Biocare Medical). Slides were then washed twice in TBS plus 0.025% Triton X-100, and endogenous peroxidase activity was blocked by a 10-minute incubation at room temperature in PeroxAbolish Solution (PXA969L Biocare Medical). Sections were then incubated for 1 hour at room temperature in the blocking solution containing the diluted secondary antibodies (HRP-conjugated goat anti-mouse IgG, 1:100, or rabbit anti-rat IgG, 1:100). Slides were washed twice in TBS, and peroxidase activity was detected with Liquid DAB Substrate Chromogen System (Dako). Nuclear counterstaining was obtained with Mayer hematoxylin. Following dehydration, sections were mounted with EUKITT (Kindler GmbH), and images acquired thanks to NIS-elements software with a Nikon Eclipse 90i digital microscope (Nikon Instruments, Europe). The percentage of positive cells was quantified using the QuPath free software for immunohistochemical analysis (Bankhead et al., 2017) as previously described (Tattersall et al., 2021).

### Statistics

All data are shown as mean ± standard error of the mean (SEM). Significance was calculated assuming equal standard deviations and variance with a two-tailed Student’s t-test, performed with the GraphPad Prism software (GraphPad, San Diego, Ca, USA). P-values lower than 0,05 were considered statistically significant.

## Results

### P2X7R deletion leads to a decrease in systemic levels of pro-inflammatory cytokines and to an increase of TGF-β

In recent years we and others have demonstrated that, in mice lacking the P2X7R, solid and liquid tumors show an increased growth due to reduced immune cell infiltration and lower eATP levels (Adinolfi et al., 2015; Hofman et al., 2015; De Marchi et al., 2019). However, an in-depth analysis of the systemic levels of several pro or anti-inflammatory cytokines in tumor-bearing P2X7R null mice has never been carried out. Figure 1 shows that, in parallel with an accelerated tumor growth (Fig.1 A-C), P2X7R null mice also show a generalized reduction of systemic pro-inflammatory cytokines (Fig.1 D-I) as well as an increase of the immune suppressant TGF-β (Fig.1 J). Notably, blood concentration of IL-6 (Fig. 1 F), IL-12 (Fig.1. G), IL-17 (Fig. 1H) and IFN-γ (Fig.1 I) was strikingly decreased (more than halved), suggesting a substantial impairment of anti-tumoral immune responses. On the contrary, blood TGF-β was almost triplicated (Fig.1 J). Taken together, these data strongly suggest that the increased tumor progression observed in P2X7R null mice could be linked to a downmodulation of the tumor-eradicating immune response.

**Figure 1:**
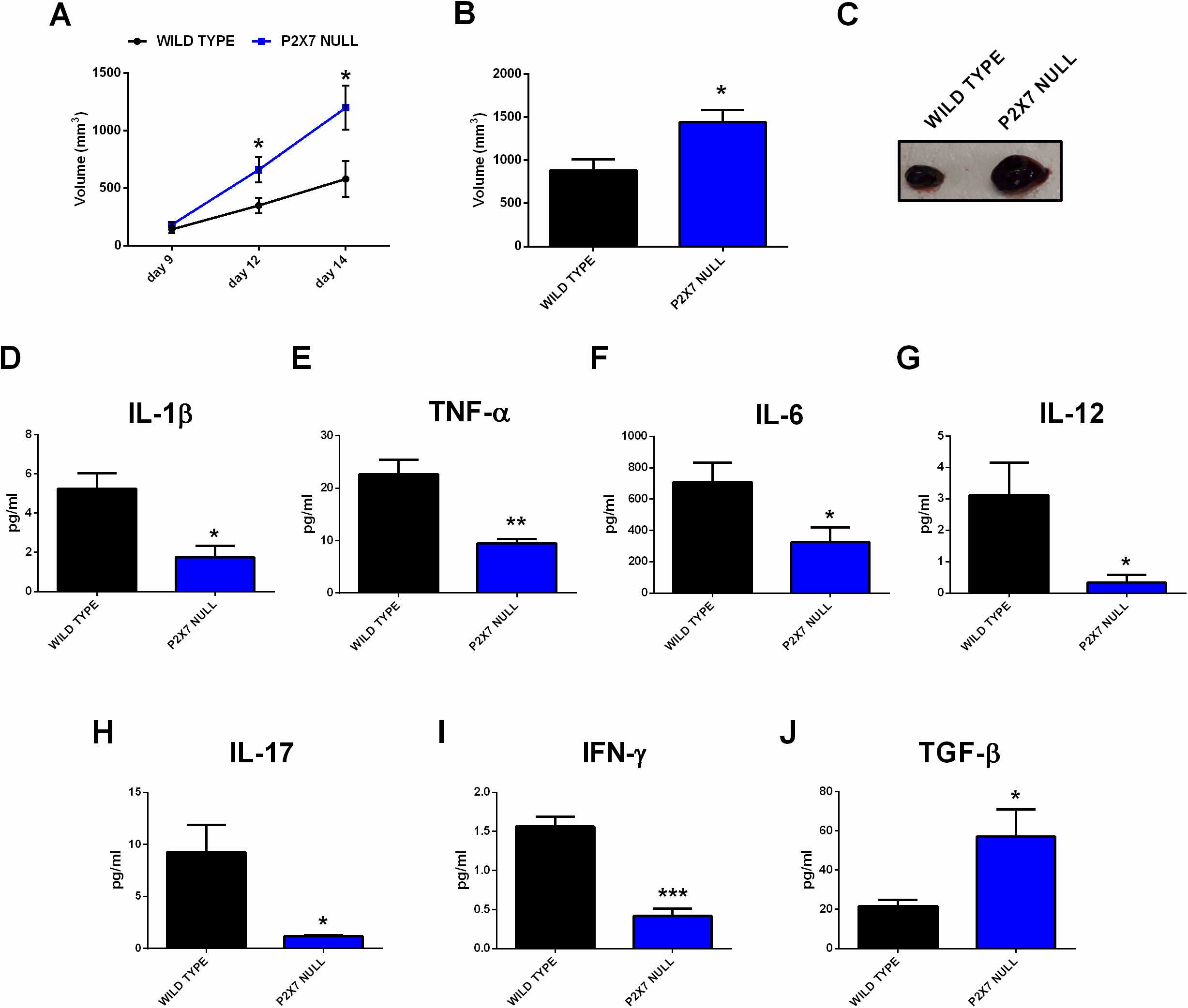
C57bl/6 mice were inoculated into the right flank with B16-F10 cells in wild type and P2X7-null mice. (**A**) Tumor volume was in vivo assessed at the indicated time points. (**B**) Ex-vivo tumor volume assessed by calliper. (**C**) Representative pictures of tumors from wild type and P2X7 null mice at post-inoculum day 14. (**D-J**) Levels of plasma cytokines of tumor-bearing C57bl/6 mice inoculated with B16 cells. IL-1β (**D**), TNF-α (**E**), IL-6 (**F**), IL-12 (**G**), IL-17 (**H**), IFN-γ (**I**) and TGF-β (**J**) were evaluated in plasma samples obtained at post-inoculum day 14. Data are shown as the mean ± SEM. *P < 0.05, **P < 0.01 and ***P < 0.001.

### The A2AR is overexpressed in tumor-bearing P2X7R null mice

Tumor immune suppression, in analogy to that observed in the P2X7R null melanoma model, has been associated with the accumulation of eADO in the TME and the ensuing activation of the A2AR (Ohta et al., 2006; Ohta and Sitkovsky, 2014). Moreover, decreased eATP concentration accompanied by increased CD73 and CD39 expression strongly support eADO accumulation in the TME of P2X7R null mice. However, P2X7R and A2AR reciprocal crosstalk in the TME was never investigated before. Therefore, we analyzed A2AR expression levels in tumors growing in either WT or P2X7 null mice observing a significant increase of tumor A2AR expression (Fig. 2 A). Interestingly, A2AR expression was also higher in the spleens, but not in the livers, from *p2x7^-/-^* mice whether in tumor-bearing or tumor free (Fig. 2 B-E).

**Figure 2:**
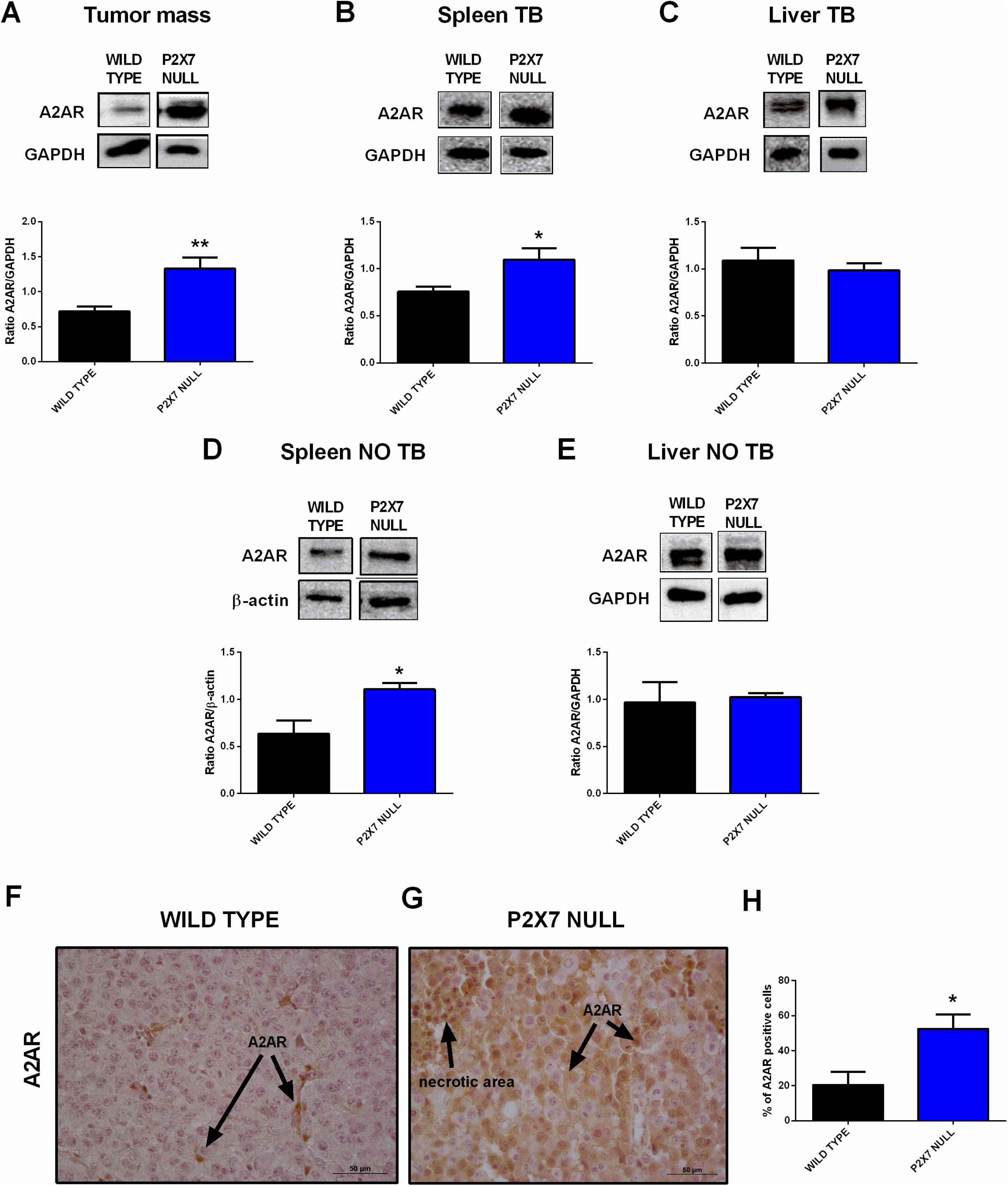
(**A-E**) Western Blot and relative quantification of A2AR in tumor (**A**), spleen (**B, D**) and liver (**C, E**) derived from wild type and P2X7 null tumor-bearing (**A-C**) or non-tumor-bearing (**D, E**) C57bl/6 mice. (**F-H**) Immunohistochemistry (**F, G**) and relative quantification (**H**) of A2AR in tumor derived from wild type (**F**) and P2X7 null (**G**) C57bl/6 mice. Pictures are obtained with a 40X objective. Data are shown as the mean ± SEM.*P<0.05 and **P<0.01.

Immunohistochemical analysis confirmed overexpression of A2AR in tumors from P2X7R-null mice (Fig. 2 F-H), mainly on tumor cells (Fig. 2 F, G) and in the periphery of necrotic sites (Fig. 3 A,B). This observation leds us to investigate tumor levels of the angiogenetic factor VEGF. VEGF is upregulated in hypoxic regions associated with necrosis, and can be secreted in response to A2AR stimulation. Figure 3 shows a significantly higher intra-mass accumulation of VEGF in tumors growing in P2X7R devoid mice as compared to WT controls (Fig. 3C). This VEGF increase was paralleled by an augmented expression of the endothelial marker CD31 (Fig.3 D-F), indicating increased intratumor blood vessel formation.

**Figure 3:**
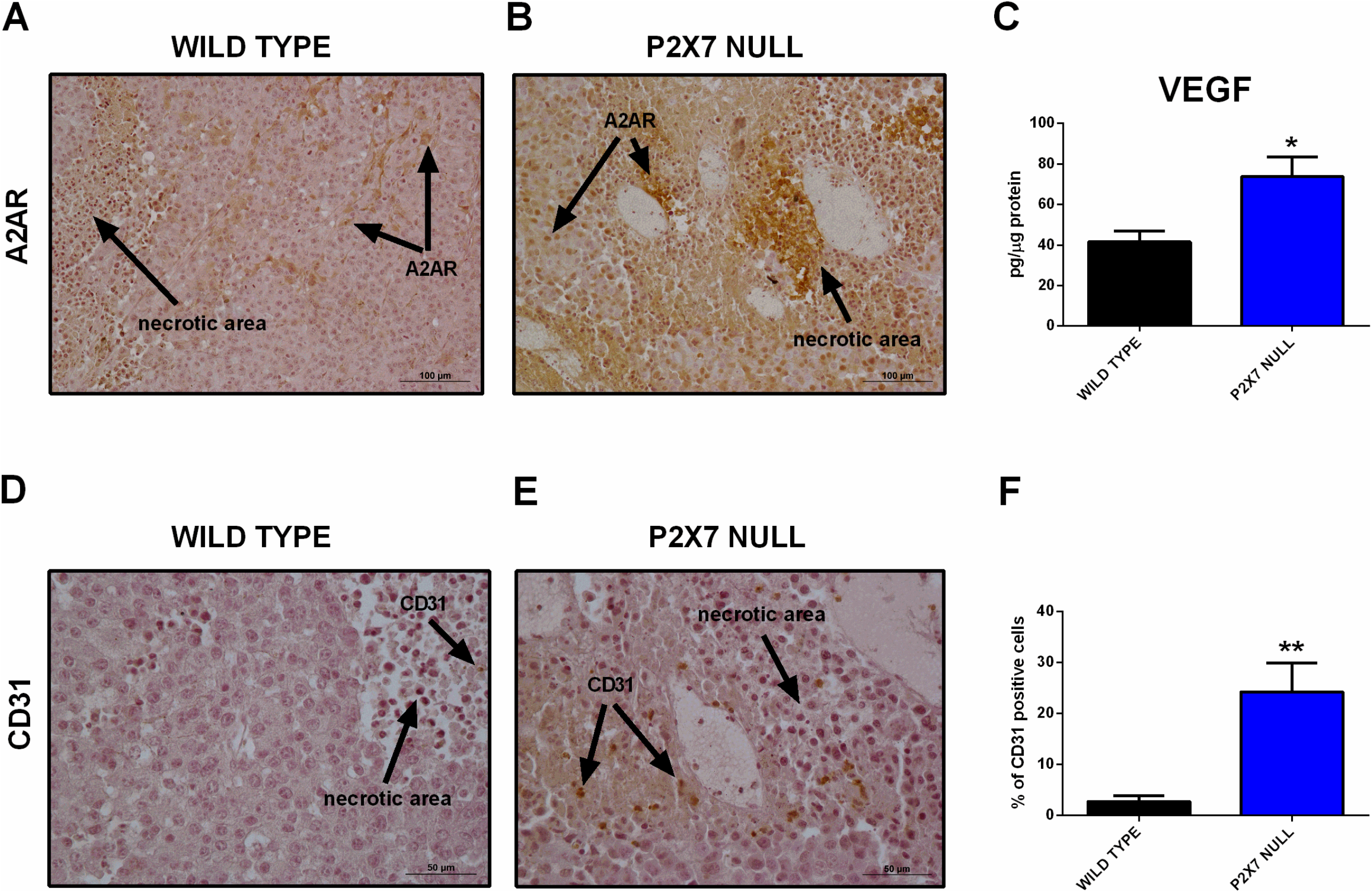
(**A, B**) Immunohistochemistry of A2AR in tumors derived from wild type (**A**) and P2X7 null (**B**) C57bl/6 mice. Pictures are obtained with a 20X objective. (**C**) Intra-tumor levels of VEGF. (**D-F**) Immunohistochemistry (**D, E**) and relative quantification (**F**) of CD31 in tumor derived from wild type and P2X7 null C57bl/6 mice. Data are shown as the mean ± SEM. *P<0.05 and **P<0.01.

### A2AR blockade reduces tumor growth in P2X7R genetically deleted mice and normalizes VEGF and TGF-β levels

To better understand the role played by A2AR in tumor progression in P2X7R-null mice, we administered the A2AR antagonist SCH58261 to *p2x7^-/-^* and WT tumor-bearing mice. This drug was administered by intraperitoneal injections at a 1 mg/kg dose every three days from the first appearance of the tumor mass. The A2AR antagonist reduced tumor growth in both mice strains (Fig. 4, A and B), thus demonstrating the central role of A2AR in the B16-F10 melanoma model. Interestingly, SCH58261 did not affect the tumor levels of A2AR, which remained significantly higher in P2X7R-null versus WT mice (Fig. 4 C,D). On the other hand, the level of P2X7R expressed by tumor cells increased in P2X7 null mice following A2AR antagonism (Fig. 4 E,F), thus further supporting an interdependency of the two receptors in the TME. It as to be noted that B16-F10 cells injected in P2X7R-null mice still express P2X7R and are, therefore, responsible for the detection of the protein shown in Fig. 4 E,F. A2AR antagonism also reduced VEGF levels only in P2X7 devoid hosts (Figure. 4 G). On the other hand, SCH58261 treatment significantly reduced TGF-β levels in tumors growing in both mice strains, further emphasizing the leading role of A2AR in generating an immune-suppressed TME.

**Figure 4:**
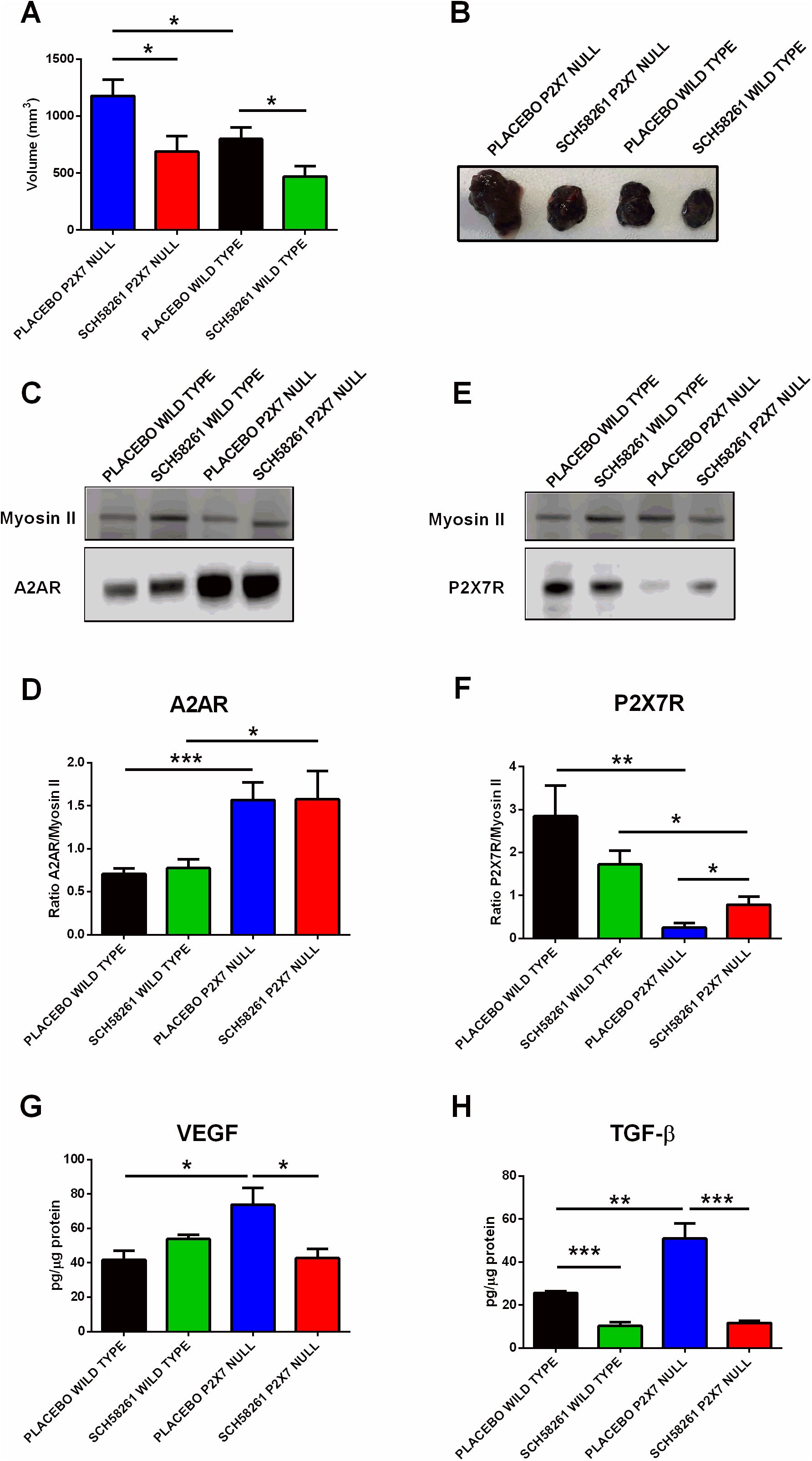
C57bl/6 mice were inoculated into the right flank with B16-F10 cells in wild type and P2X7 null mice. A2AR antagonist SCH-58261 (1 mg/kg) or placebo (sterile PBS containing 0.005% DMSO) were intra-peritoneum injected every 3 days after first tumor mass detection (day 5 from inoculum). (**A**) Ex-vivo tumor volume assessed by calliper. (**B**) Representative pictures of tumors derived from treated wild type and P2X7 null mice at post inoculum day 14. (**C-F**) A2AR (C) and P2X7R (**E**) expression by Western Blot and relative quantification (**D, F**). VEGF (**G**) and TGF-β (**H**) were evaluated in tumor masses obtained at post inoculum day 14. Data are shown as the mean ± SEM. *P < 0.05, **P < 0.01 and ***P < 0.001.

## Discussion

The TME is the privileged site of host-tumor interaction and, as such, a key determinant of cancer progression and metastatic spreading. Properties of the TME are also dictated by its features, such as hypoxia that can determine tumor vascularization causing the release of VEGF. Over the last few years, the abundance of eATP was unexpectedly identified as a prominent TME characteristic (Di Virgilio et al., 2018). eATP accumulates into the TME as a consequence of cell death or injury and of non-lytic release from tumor and host cells (Di Virgilio et al., 2018). In the TME, eATP ligates its specific cognate receptors (P2Y and P2X) and is metabolised by plasma membrane-expressed or soluble ecto-ATPases. The semi-final degradation product of eATP is eADO (the final being inosine), a potent immunosuppressant acting at A2AR (Sitkovsky et al., 2014; Vijayan et al., 2017; Boison and Yegutkin, 2019). Thus, eATP is central in the immunoregulation of the TME because, on the one hand, it is an immunostimulant acting at P2Y and P2X receptors, and on the other hand supports immunosuppression by generating eADO and activating A2AR. Converging evidence supports the view that the P2X7R is the P2 receptor most heavily involved in host-tumor interaction in the TME (Lara et al., 2020). eATP and the P2X7R are “partners in crime” in the TME as eATP ligates P2X7R and triggers P2X7R-mediated responses, among which release of ATP-itself, controlling the concentration of its own agonist (De Marchi et al., 2019). We recently demonstrated a substantial reduction in eATP levels in tumor-bearing mice devoid of P2X7R. This reduction was also accompanied by a significant increase of ectonucleotidases expressed by tumor-infiltrating immune cells, in particular Tregs, thus suggesting an upregulation of eADO levels in the TME of mice lacking P2X7R (De Marchi et al., 2019).

In the current study, we completed these observations by concentrating on another key actor of the adenosinergic pathway in cancer: the A2AR (Sitkovsky, 2020). Indeed, A2AR role in causing tumor immune suppression in hypoxic tumors is a consolidated notion that recently lead to several clinical trials exploring the anti-cancer therapeutic efficacy of the administration of A2AR antagonists as a stand-alone treatment or in combination with immune checkpoints inhibitors (Boison and Yegutkin, 2019; Fong et al., 2020; Franco et al., 2021; Augustin et al., 2022). Interestingly, in the TME A2AR activation has been reported to reduce the production of those tumor eradicating cytokines IL-6, IL-17 and IFN-*γ* that we fund downmodulated in tumor-bearing *p2x7^-/-^* mice in the present study (Augustin et al., 2022). We also observed a reduction of other pro-inflammatory cytokines (IL-1β, TNF-α and IL-12) related to P2X7R activation and, in general, to innate and type 1 immune response (Correa et al., 2017; Orioli et al., 2017). This immune, cancer-promoting scenario prompted us to further analyse A2AR levels in B16-F10 melanoma-bearing *p2x7^-/-^* and WT mice model. Our data show that in P2X7-null mice A2AR expression was increased in the tumor cells and in immunocompetent tissues such as the spleen. However, A2AR expression was also increased in the spleen of non-tumor-bearing P2X7-null mice, strongly suggesting that P2X7R deletion is a prime driver of A2AR upregulation in the immune compartment. In support of this hypothesis, no increased A2AR expression was found in liver and other non-immune tissues from *p2x7^-/-^* mice (not shown). Of interest, A2AR inhibition was able to increase P2X7R expression by B16-F10 cells, thus suggesting that the compensatory mechanism leading to upregulation of one of the two receptors when the other is downmodulated or blocked might be reciprocal.

A2AR is also known to promote neovascularisation and immune suppression via VEGF and TGF-β release (Steingold and Hatfield, 2020) both upregulated in melanoma-bearing *p2x7^-/-^* mice. Moreover, to further stress the pivotal function of the eADO signalling in the tumor hypoxic response, A2AR expression in the *p2x7^-/-^* mice was enhanced in peri-necrotic areas, together with an enhanced staining by the endothelial marker CD31. Of note, A2AR inhibition normalised intra-tumor VEGF levels only in P2X7-null mice, thus further supporting the hypothesis that the pro-angiogenic phenotype observed in this model was mainly dependent upon A2AR activity. Finally, A2AR blockade inhibited TGF-β release, thus further reducing immune suppression in the TME.

Our data unveil a new A2AR dependent compensatory mechanism emerging in the absence of host P2X7R that favours tumor growth via immune suppression and neovascularisation. Moreover, one should consider that when targeting P2X7R within cancer, it might also influence A2AR activity and vice-versa and, thus, that an anti-tumoral therapeutic strategy targeting both receptors has the potential to prove more efficacious than a stand-alone one. Finally, based on previous studies, a combination of P2X7R and A2AR directed drugs could also prove beneficial in association with traditional chemotherapy (Pegoraro et al., 2020; Sitkovsky, 2020) or irradiation (Huang et al., 2020; Zanoni et al., 2022).

## Author Contributions

EDM performed most of the experimental work. AP helped with in vivo and immunohistochemistry experiments. RT helped with Western blotting analysis. FDV, SM, EDM, AP participated in experimental design and manuscript preparation. EA designed and supervised the study, participated in in vivo experimentation, wrote and gave final approval to the manuscript.

## Acknowledgements

The authors are grateful to Marzia Scarletti and Paola Chiozzi for technical support, and to Anna Scialdone and Letizia Alfieri for fruitful discussion.

## Data Availability Statement

Complete raw data sets presented in this study will be made available upon reasonable request to the corresponding author.

